# Functional analysis of cell lines derived from SMAD3-related Loeys-Dietz Syndrome patients provides insights into genotype-phenotype relations

**DOI:** 10.1101/2023.12.11.571192

**Authors:** Nathalie P. de Wagenaar, Lisa M. van den Bersselaar, Hanny J.H.M. Odijk, Sanne J.M. Stefens, Dieter P. Reinhardt, Jolien W. Roos-Hesselink, Roland Kanaar, Judith M.A. Verhagen, Hennie T. Brüggenwirth, Ingrid M.B.H. van de Laar, Ingrid van der Pluijm, Jeroen Essers

**Author notes:** Corresponding authors, Ingrid M.B.H. van de Laar, Department of Clinical Genetics, Erasmus University Medical Center, P.O. Box 2040, 3000 CA Rotterdam, The Netherlands. Mail address; Ingrid van der Pluijm, Erasmus University Medical Center Rotterdam, Room Ee702b, Wytemaweg 80, 3015 CN Rotterdam, The Netherlands. Mail address, Jeroen Essers, Erasmus University Medical Center Rotterdam, Room Ee702b, Wytemaweg 80, 3015 CN Rotterdam, The Netherlands. Mail address. These authors share first authorship (Nathalie P. de Wagenaar, Lisa M. van den Bersselaar).

## Abstract

**Introduction:** Pathogenic (P) and likely pathogenic (LP) variants in the *SMAD3* gene cause Loeys-Dietz syndrome type 3 (LDS3), also known as aneurysms-osteoarthritis syndrome (AOS). The phenotype of LDS3 is highly variable and characterized by arterial aneurysms, dissections and tortuosity throughout the vascular system combined with skeletal, cutaneous and facial features.

**Objectives:** Investigate the impact of P/LP *SMAD3* variants through conducting functional tests on patient-derived fibroblasts and vascular smooth muscle cells (VSMCs).The resulting knowledge will optimize interpretation of *SMAD3* variants.

**Material and methods:** We conducted a retrospective analysis on clinical data from individuals with a P/LP *SMAD3* variant and utilized patient-derived VSMCs to investigate the functional impacts of dominant negative (DN) and haploinsufficient (HI) variants in SMAD3. Additionally, to broaden our cell model accessibility, we performed similar functional analyses on patient-derived fibroblasts carrying SMAD3 variants, differentiating them into myofibroblasts with the same variants. This enabled us to study the functional effects of DN and HI variants in *SMAD3* across both patient-derived myofibroblasts and VSMCs.

**Results:** Individuals with dominant negative (DN) variants in the MH2 protein interaction domain of SMAD3 exhibited a higher frequency of major events (66.7% vs. 44.0%, p=0.054), occurring at a younger age compared to those with haploinsufficient (HI) variants. Moreover, the age at the onset of the first major event was notably younger in individuals with DN variants in MH2, 35.0 years [IQR 29.0-47.0], compared to 46.0 years [IQR 40.0-54.0] in those with HI variants (p=0.065). In functional assays, fibroblasts carrying DN *SMAD3* variants displayed reduced differentiation potential, contrasting with increased differentiation potential observed in fibroblasts with HI *SMAD3* variants. Additionally, HI *SMAD3* variant VSMCs showed elevated SMA expression, while exhibiting altered expression of alternative MYH11 isoforms. Conversely, DN *SMAD3* variant myofibroblasts demonstrated reduced extracellular matrix (ECM) formation compared to control cell lines. These findings collectively indicate distinct functional consequences between DN and HI variants in *SMAD3* across fibroblasts and VSMCs, potentially contributing to the observed differences in disease manifestation and age of onset of major events.

**Conclusion:** Distinguishing between P/LP HI and DN *SMAD3* variants can be achieved by assessing differentiation potential, and evaluating SMA and MYH11 expression. Notably, myofibroblast differentiation seems to be a suitable alternative in vitro test system in comparison to VSMCs. Moreover, there is a notable trend of aortic events occurring at younger age in individuals with a DN *SMAD3* variant in the MH2 domain, distinguishing them from those with a DN variant in the MH1 domain or a HI variant.

## INTRODUCTION

An aortic aneurysm is a local widening of the aorta (1). Increased aortic diameter is directly associated with an increased risk of aortic dissection or rupture, a life-threatening event (2). Several genes are associated with aortic aneurysm formation, including genes affecting the TGF-β pathway, the extracellular matrix (ECM), and smooth muscle cell contraction (3).

Genes involved in the TGF-β signaling pathway related to aneurysm formation include *TGFBR1* (LDS1), *TGFBR2* (LDS2), *SMAD3* (LDS3), *TGFB2* (LDS4), *TGFB3* (LDS5), *SMAD2* (LDS6), *SMAD4* (juvenile polyposis/hereditary hemorrhagic telangiectasia), *SMAD6* (bicuspid aortic valve/thoracic aortic aneurysm) and *SKI* (Shprintzen-Goldberg syndrome) (4–14). *SMAD3* is a signal transducer in the canonical TGF-β pathway. Upon activation by binding of TGF-β, the TGF-β receptor complex activates SMAD3 and SMAD2 proteins (receptor-regulated Smads; R-SMADs) by phosphorylation. Once phosphorylated, the R-SMADs can form a protein complex with SMAD4 (co-SMAD). Those complexes then transfer to the nucleus where they interact with promotors/co-factors to regulate the transcription of downstream genes, including genes involved in ECM turnover, apoptosis and cellular proliferation, differentiation, motility and adhesion (15–22).

The SMAD3 protein consists of two functional domains separated by a linker region; the N-terminal domain Mad Homology 1 (MH1) and C-terminal domain Mad Homology 2 (MH2). MH1 enables DNA binding and MH2 mediates protein-protein interaction and SMAD-dependent downstream transcription (23). No mutational hotspots are identified in *SMAD3*, but the majority of the pathogenic/likely pathogenic (P/LP) missense variants are located in the MH2 domain (24).

Heterozygous P/LP variants in *SMAD3* (OMIM 603109) cause Loeys-Dietz Syndrome type 3 (LDS3, OMIM 613795, ORPHA 284984), also known as aneurysms-osteoarthritis syndrome (AOS). LDS3 is characterized by aneurysms and tortuosity of the aorta and/or middle-sized arteries, accompanied by osteoarthritis (5, 6). Currently, over 60 different P/LP variants in *SMAD3* have been identified in LDS3 families (24–28), including missense, truncating and splicing variants, and intragenic and whole gene deletions (24, 29, 30). Interestingly, large phenotypic variation is described between LDS3 families, suggesting that, among others, genotype and ancestry might play a role. Hostetler et al. showed that an aortic event occurred at younger age in individuals with a DN variant in the MH2 domain of SMAD3 compared to individuals with a HI variant in the MH2 domain or a variant in the MH1 domain (31). In addition, variability in severity of the phenotype is observed within families (5, 32), which indicates that the severity of LDS3 is influenced by other factors, including genetic modifiers (33), lifestyle, co-morbidities, and/or sex.

Besides P/LP *SMAD3* variants, variants of unknown significance (VUS) are often found in *SMAD3*. In our centre, 13 out of 50 unique *SMAD3* variants (26%) discovered upon diagnostic testing were classified as VUS, including 10 variants found in syndromic TAA patients and 3 variants in non-syndromic TAA patients (unpublished data). Improving the interpretation of the pathogenicity of these *SMAD3* VUS is essential, since the molecular genetic diagnosis is important for appropriate management and treatment of the patient and allows for predictive genetic testing in family members at risk. A molecular genetic diagnosis of LDS3 guides the frequency and extent of vascular imaging and the threshold for preventive surgery (34). The vast amount of VUS found in genetic testing as well as the clinical utility of a certain diagnosis of LDS3, underlines the need for assays to functionally characterize and interpret *SMAD3* variants.

Here, we studied the effect of P/LP and VUS *SMAD3* variants by performing functional experiments on patient-derived in vitro differentiated fibroblasts (myofibroblasts) and VSMCs. Myofibroblasts differentiation can be a more accessible cell model to study patient-related variants compared to VSMCs, since VSMCs can only be obtained during surgery. In these functional cell-based experiments, we analyzed the differentiation potential, activation of the TGF-β pathway, the expression of SMC markers, and the formation of ECM.

## MATERIAL AND METHODS

### Patient cells and characteristics

Primary dermal fibroblasts and VSMCs were collected from individuals with a P/LP variant or VUS in the *SMAD3* gene (NM_005902.4). In total, five *SMAD3* patient fibroblast cell lines, two control fibroblast cell lines (Biobank Clinical Genetics, Erasmus MC), seven *SMAD3* patient VSMC lines and three commercially available control VSMC lines were used (Lonza CC-2571, lot no. 0000369150, ATTC PCS-100-012, lot no. 62726859 and lot no. 64193202) (Supplementary table 1). Collection of patient material was approved by the medical ethics committee of the Erasmus Medical Center Rotterdam (MEC 2014-579 and MEC 2017-040) and written informed consent was provided by all patients. A retrospective analysis was performed to obtain clinical data from the 12 individuals of the included samples (Supplementary table 2). In addition, we reviewed the clinical and genetic data of all LDS3 67 individuals clinically examined in Rotterdam with a confirmed P/LP *SMAD3* variant, using the same methodology (Table 3, Supplementary table 5).

### Cell culture

VSMCs were cultured in SmGM-2 medium supplemented with SMBM growth factors (Lonza, CC-4149 and CC-3181) in gelatin-coated dishes and incubated at 37°C with 5% CO_2_. Fibroblasts were cultured in Dulbecco’s Modified Eagle’s Medium (DMEM, Gibco, #11965-092) supplemented with 10% fetal calf serum (FCS, Capricorn, FBS-12A) and 1% penicillin/streptomycin (P/S, Sigma, P0781). The cultured fibroblasts were incubated at 37°C with 5% CO_2_.

### RNA isolation and cDNA

RNA was isolated from VSMCs with the RNeasy mini kit (Qiagen, #74104). cDNA was prepared with the iScript cDNA synthesis kit (Biorad, #170-8882) according to the manufacturer’s protocol.

### Restriction analysis

Restriction enzyme cleavage of PCR products derived from cDNA was performed to confirm the P/LP variants in these cell lines (Table 1). PCR on cDNA was performed with Q5 polymerase according to the manufacturer’s protocol (Biolabs, M0493S). The PCR product was digested with an enzyme that gains or loses a restriction site due to the specific P/LP variant in that fragment (Table 1, NEB). Digestion was performed according to the manufacturer’s protocol. After restriction, the products were separated on 2% agarose gels.

**Table 1.**
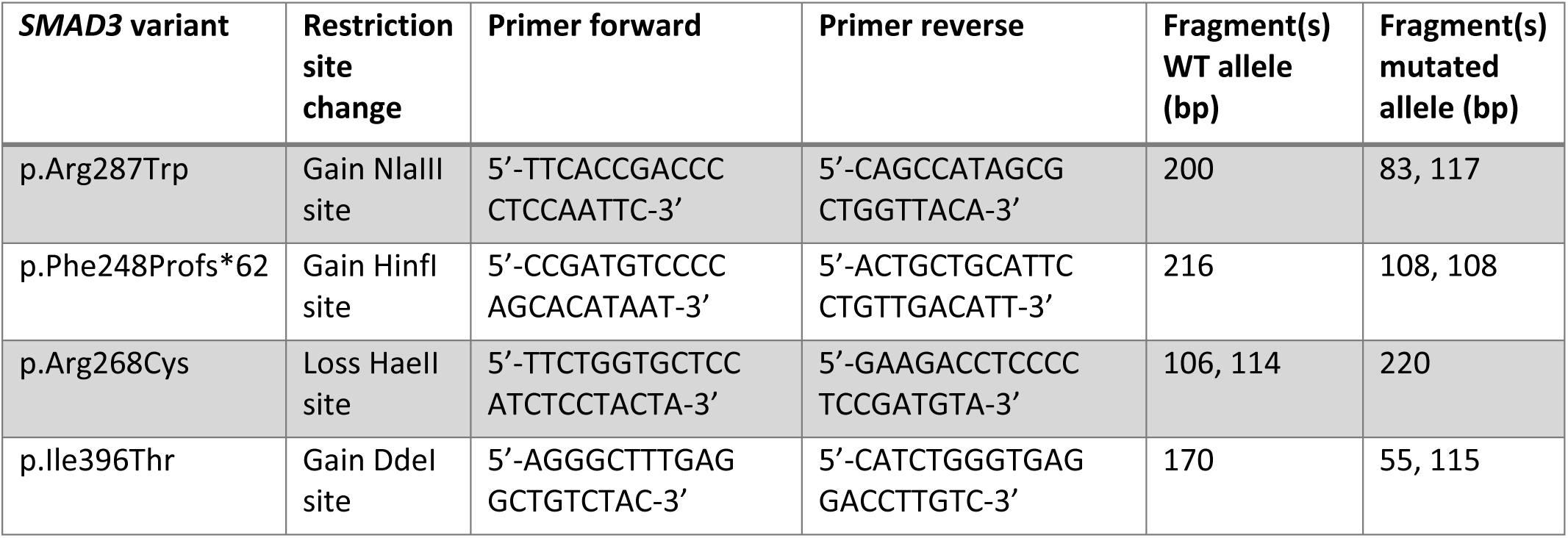
Primers and expected sizes restriction analysis *SMAD3*.

### TGF-β stimulation

Cells were seeded in 6-well plates with DMEM, 1% P/S and 10% FCS to reach confluence. After 24 hours, the medium was replaced by FCS-free medium. Cells were then stimulated with TGF-β for 0 minutes, 15 minutes, 30 minutes, 1 hour and 4 hours before collecting protein lysates. Lysis and western blotting were performed as described in the Western blotting protocol below.

### Differentiation into myofibroblasts

Patient and control fibroblasts were differentiated into myofibroblasts as published (35) (Figure 3A). In short, 250.000 fibroblasts were seeded in a 1 cm^2^ piece of Matriderm (MedSkin Solutions, 83403-200). After 2 days, cells were attached and medium was replaced by DMEM with 2% heat-inactivated FCS, 1% P/S and 5ng/ml active human recombinant TGF-β1 (Biovision, 4342-5). This medium was replaced every 4 days. At day 14, cells were enzymatically extracted from Matriderm by adding collagenase (2000 IU/ml, Worthington, LS004176) in DMEM with 10% FCS and 1% P/S and shaking at 37°C for 3 hours. A centrifugation step was performed to remove collagenase before the myofibroblasts were seeded in new flasks and cultured in DMEM with 10% FCS and 1% P/S. After completion of the differentiation protocol, cells were used after 2-3 passages for functional assays. Passage number was kept below 5 to prevent loss of differentiation markers.

### Western blotting

VSMCs and myofibroblasts were scraped in phosphate-buffered saline (PBS) supplemented with protease inhibitor cocktail (1:100, Roche, 11836145001) and phosphatase inhibitor cocktail (1:100, Sigma, P0044). These samples were lysed in equal volumes of 2x Laemmli buffer (4% SDS, 20% glycerol, 120mM Tris pH 6,8) supplemented with protease inhibitor cocktail and phosphatase inhibitor. Lysates were passed through a 25G needle and heated to 65°C for 10 minutes. Protein concentrations were measured with Lowry protein assay (36). Equal amounts of protein were separated on a SDS-PAGE gel and transferred to an Immobilon-P polyvinylidene difluoride (PDVF) membrane (Millipore, IPVH00010). Membranes were blocked in PBS with 3% milk powder (Sigma, 70166) and 0.1% Tween-20 (Sigma, P1379). After blocking, the membranes were incubated overnight in blocking buffer with primary antibody (Supplementary table 3). Membranes were washed with 0,1% Tween-20 in PBS (5 times) and incubated for 1 hour at room temperature (RT) with horseradish peroxidase-conjugated secondary antibodies (1:2000, Jackson ImmunoResearch, 515-035-003 and 711-035-152). Membranes were washed again before protein detection, using home-made enhanced chemiluminescence (ECL) substrate on Amersham Imager 600 (GE Healthcare Life Sciences).

Quantification of protein signals was performed using Fiji software (37).

### Immunofluorescent staining

Subconfluent VSMCs (50-70%) were grown on 18 mm coverslips in 12-well plates and fixed with 4% paraformaldehyde in PBS for 30 minutes. For immunostaining of differentiated fibroblasts, 100,000 skin fibroblasts/well were seeded on 18 mm coverslips in 12-well plates in DMEM with 10% FCS and 1% PS. After 2 days, medium was replaced by DMEM supplemented with 2% heat-inactivated FCS, 1% PS and 5ng/ml human recombinant TGF-β1 (Biovision, 4342-5). This medium was changed every 4 days and after 14 days, cells were fixed with 2% paraformaldehyde in PBS for 15 minutes. After fixation, cells were permeabilized with PBS supplemented with 0.1% Triton-X-100 and blocked with PBS+ (PBS with 0.5% bovine serum albumin (BSA) and 0.15% glycine) for 30 minutes. Coverslips were incubated overnight at 4°C in PBS+ with primary antibodies: mouse monoclonal anti-smooth muscle actin (SMA) (1:750, Abcam, ab7817) and rabbit polyclonal anti-SM22 (1:400, Abcam, ab14106) or rabbit polyclonal anti-MYH11 (1:500, Abcam, ab53219) or rabbit monoclonal anti-vimentin (1:1000, Abcam, ab92547). Coverslips were washed with PBS supplemented with 0.1% Triton-X-100 and shortly with PBS+ before incubation with the secondary antibodies; anti-mouse Alexa Fluor 488 (1:1000, Molecular Probes, A11001) and anti-rabbit Alexa Fluor 594 (1:1000, Molecular Probes, A11012) and SiR-actin probe (1:1000, Cytoskeleton, SC001) in PBS+ for 1 hour at room temperature. Coverslips were mounted in Vectashield with Dapi (Vector laboratories, H-1200,) and sealed with nail polish. Images were recorded with an Axio Imager D2 microscope (Zeiss).

### Immunofluorescence of extracellular matrix

Cells were seeded at 50,000 cells/cm^2^ in 8-well removable chamber slides (Ibidi, 80841) and grown for 7 days to allow ECM deposition. Cells were fixed with ice-cold 70:30 methanol: acetone mixture for 5 minutes and washed with PBS. Blocking was performed for 1 hour in PBS supplemented with 10% normal goat serum (NGS) (Agilent, X0907). The cells were incubated overnight at 4°C with primary antibodies (Supplementary table 4) in PBS + 10% NGS. After washing with PBS + 0.05%Triton X-100 (3 times for 5 minutes), the coverslips were incubated with a secondary antibody in PBS + 10% NGS for 1.5 hours at ambient temperature (Molecular Probes, anti-rabbit Alexa Fluor 594, 1:1000, A11012). Coverslips were washed with PBS + 0.05% Triton X-100 and mounted to glass slides with Vectashield with DAPI (Vector laboratories, H-1200) and sealed with nail polish. Images were recorded with an Axio Imager D2 microscope (Zeiss).

### PCR *MYH11* isoforms

Primers were designed for the different *MYH11* isoforms (Table 2). PCR on cDNA was performed with Q5 polymerase (NEB, M0491) according to the manufacturer’s protocol. PCR products were separated on 2% agarose gels.

**Table 2.**
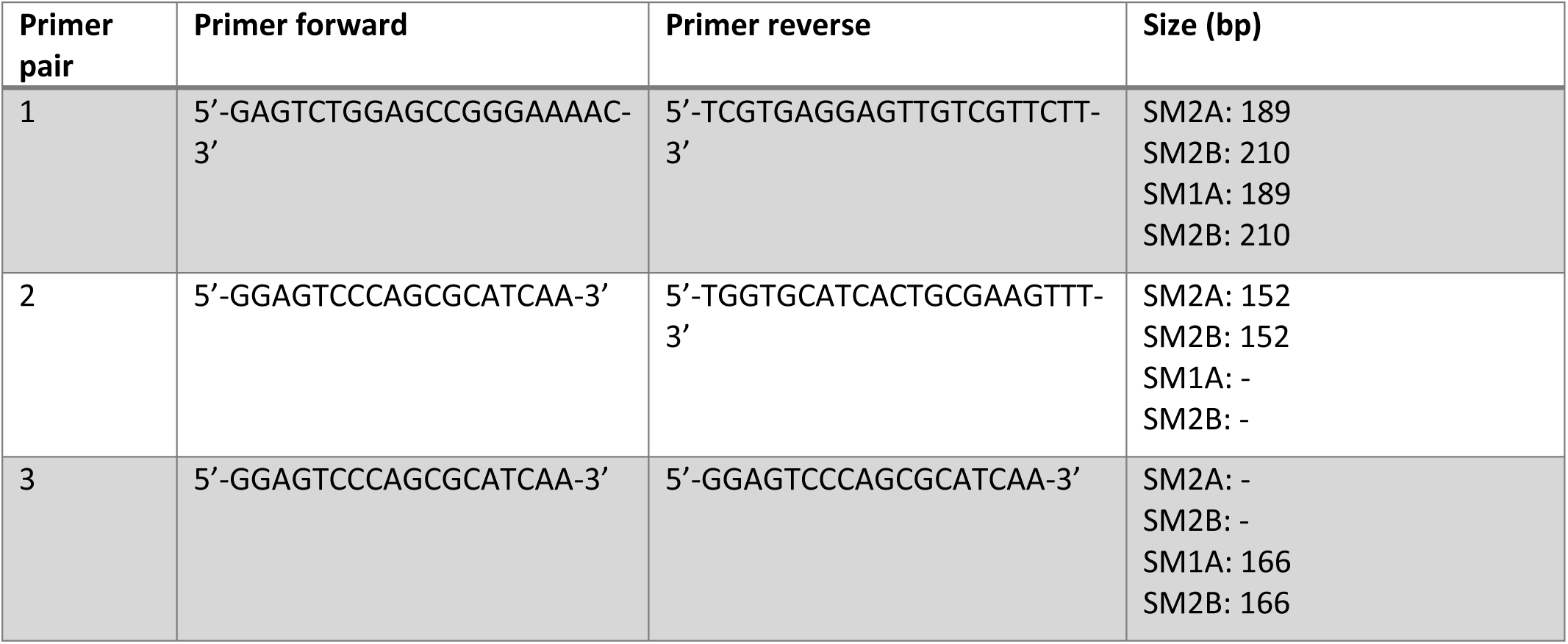
Primer pairs and expected sizes MYH11 isoforms.

### RNA sequencing and splice analysis

RNA was isolated from VSMCs as described above. Library preparation, sequencing and primary data analysis were performed at GenomeScan B.V. (Genomescan B.V., Leiden, The Netherlands). Libraries were paired-end sequenced in an Illumina Novaseq6000 platform at a sequencing depth of 40 million reads and 150 bp read length. Reads were aligned with the reference genome GRCh37.p13. To investigate alternative splicing of *MYH11*, sashimi plots were created in Integrative Genomics Viewer (IGV) (38). Minimal junction coverage was set to two.

### Statistics

Data were corrected for outliers with the Grubbs’ test for outliers. Statistical analysis was performed with a non-parametric Mann-Whitney test. For a significant difference between groups the p-value must be <0.05. In addition, the 1x standard deviation (SD) value of the controls is shown in the figures. All analyses were performed using Graphpad Prism, version 8.

## RESULTS

### Phenotype-genotype correlations

In our center, we identified 67 individuals with LDS3 from 12 families (Table 3 and Supplementary table 5). Over half of the individuals (41/67, 61.2%) were heterozygous for a missense variant in the MH2 domain of *SMAD3*. Only one missense variant in the MH1 domain was reported (Table 3, Supplementary table 5). In total, 37/67 individuals (55.2%) suffered from an aortic event, with an overall median age of 40.5 years [IQR 31.0-50.8]. The overall median age of individuals without an aortic event was 43.5 years [IQR 21.0-60.8]. Individuals with a missense variant in the MH2 domain of SMAD3 were more likely to experience an aortic event compared to individuals with an HI variant (66.7% versus 44.0% respectively, p=0.054). The median age at aortic event for individuals with a missense variant in MH2 was 35.0 years [IQR 29.0-47.0], which, although not significantly different, was 11 years lower than the median age at aortic event for individuals with an HI variant (46.0 years [IQR 40.0-54.0], p=0.07).

**Table 3.** Genotype-phenotype correlations.

|  | **Missense MH1**<br>**domain (n=1)** | **Missense MH2**<br>**domain (n=41)** | **Haploinsufficient (n=25)** | **MH2 vs HI**<br>**P-value** |
| --- | --- | --- | --- | --- |
| **Median age at aortic event**<br>**(years)** | - | 35.0 [29.0-47.0] | 46.0 [IQR 40.0-54.0] | 0.065 |
| **Aortic event (% of total)**<br>**(n=37)** | 0 | 26 (66.7%) | 11 (44.0%) | 0.054 |
| **Median age no aortic**<br>**event (years)** | 75 | 48.5 [28.25-63.75] | 34.0 [19.5-54.5] | 0.084 |
| **Sudden death (% of total)**<br>**(n=17)** | 0 | 8 (20.5%) | 9 (36.0%) | 0.171 |
| **Osteoarthritis (% of total)**<br>**(n=36)** | 0 | 28 (71.8%) | 8 (32.0%) | 0.005 |

### Effect of *SMAD3* HI and DN variants on the protein level/structure

The *SMAD3* variants and cell lines studied are depicted in Supplementary table 1. Variant p.Phe248Profs*62 (FB#3) and the deletion of a large part of *SMAD3* (VSMC#7) are expected to result in HI (6). Based on the protein structure of the SMAD3/SMAD4 complex, we expected *SMAD3* variant p.Arg287Trp (VSMC#1-6 and FB#1-2) and p.Arg268Cys (FB#4) to have a DN effect since the change in the SMAD3 protein structure will affect the stability and formation of the SMAD3-SMAD3-SMAD4 protein complex. The interruption of the protein complex is due to a loss of positive charge, the bulky side chain of tryptophan, and potential loss of H-bridges at the interaction site (Figure 1A). The variant p.Ile396Thr (FB#5) was classified as VUS and is expected to have a DN effect. This variant is not close to the interaction site and is expected to reduce the stability of the SMAD3 protein (Figure 1A). DNA restriction analysis confirmed the expected mutant allele in all cell lines with a *SMAD3* missense variant (Figure 1B-C). No mutant allele is present in the haploinsufficient cell line FB#3 p.Phe248Profs*62, since the expression of the mutant allele will be very low or absent due to nonsense-mediated decay (NMD) (6).

**Figure 1.**
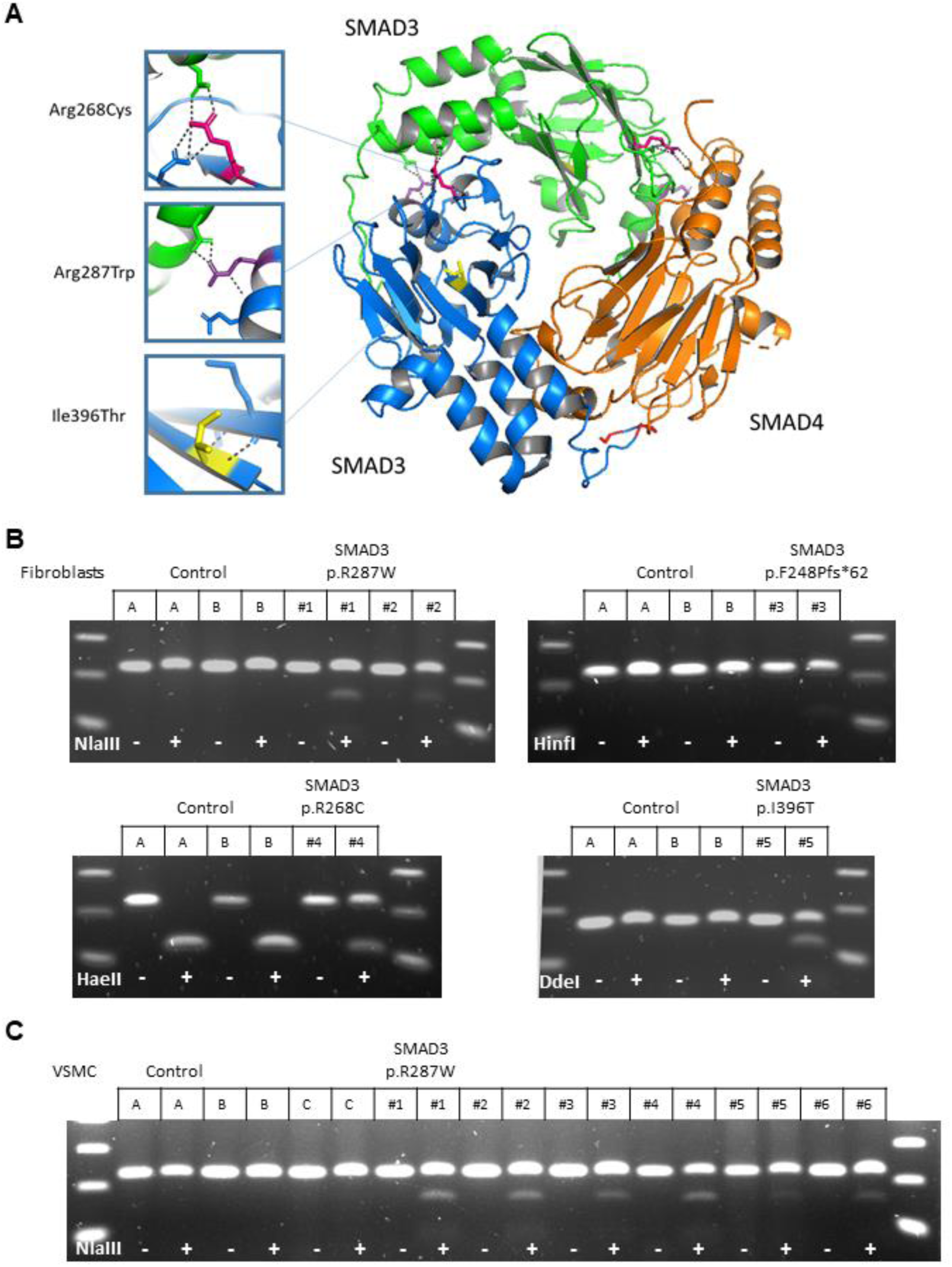
SMAD3 mutations. **A)** Protein structure of two MH2 domains of SMAD3 (blue and green) which form a protein complex together with a MH2 domain of SMAD4 (orange). Locations of DN SMAD3 mutations in patient cell lines are indicated. p.Arg287Trp (purple) and p.Arg268Cys (pink) will change the charge of the amino acid and are located near the interaction side, which will likely affect interactions between proteins and stability of the protein complex. p.Ile396Thr (yellow) will probably affect stability of protein structure. Image of PDB file 1U7F (54) created with PyMol (55). **B)** Restriction analysis of PCR product on cDNA fibroblasts. **C)** Restriction analysis of PCR product on cDNA VSMCs.

### *TGF-β* induced differentiation and functional analysis of patient-derived fibroblasts

#### Initial activation of the TGF-β pathway in SMAD3 fibroblasts is not affected

TGF-β signaling activity was determined by performing western blots for SMAD2, pSMAD2, SMAD3 and pSMAD3 at different time points after TGF-β stimulation (Figure 2). All cell lines showed induction of SMAD2 and SMAD3 phosphorylation after TGF-β stimulation. Phosphorylated SMAD2 and SMAD3 reached their highest levels 30-60 minutes after TGF-β stimulation. These data indicate that initial activation of the TGF-β pathway can still occur in cell lines with *SMAD3* variants. As expected, FB#3 p.F248Pfs*62 showed reduced expression of SMAD3 (Figure 2D). Although pSMAD3/SMAD3 ratios were unaltered in this HI cell line, the results do show that the absolute SMAD3 and pSMAD3 protein levels were lower. Due to NMD, this reduction is expected for HI variants and could lead to a lower downstream TGF-β signaling response.

**Figure 2.**
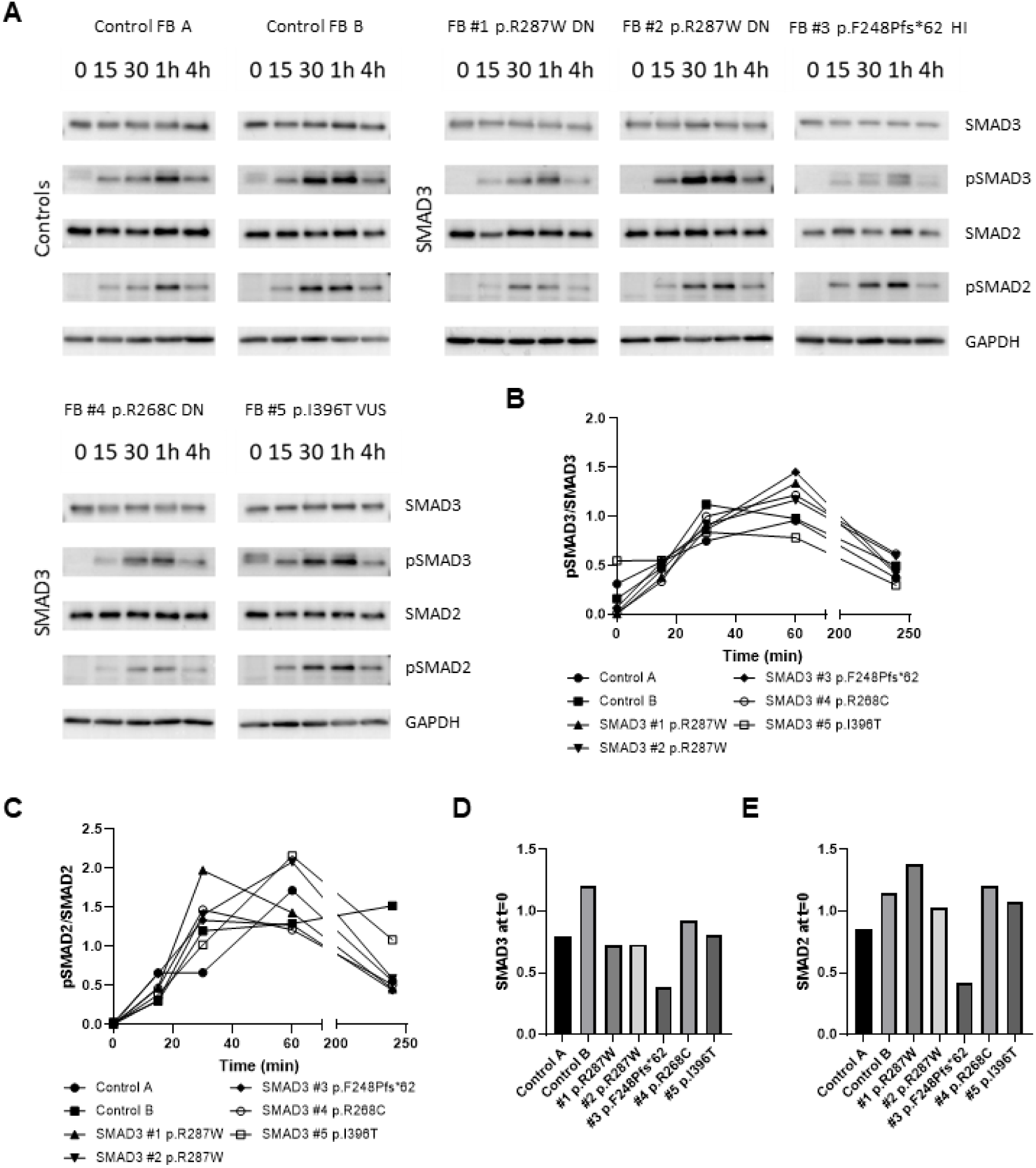
Downstream transcriptional activation of TGF-β signaling in SMAD3 fibroblasts. **A)** Western blots detecting pSMAD3, SMAD3, pSMAD2, and SMAD2 in fibroblasts upon stimulation with TGF-β (15 min, 30 min, 1 hour, and 4 hours) after serum deprivation. GAPDH levels serve as a loading control. **B)** Quantification of pSMAD3/SMAD3 ratio. **C)** Quantification of pSMAD2/SMAD2 ratio. **D)** SMAD3 levels at t=0. **E)** SMAD2 levels at t=0.

#### Reduced differentiation potential of fibroblasts with DN SMAD3 P/LP variants

During patient fibroblast culture, the morphology of *SMAD3* patient fibroblasts appeared to differ from controls (Supplementary figure 1). The *SMAD3* patient fibroblasts have a rounded morphology, while control fibroblasts have a more continuous and stretched morphology. The differences in morphology between *SMAD3* patient cells and controls were still present after differentiation into myofibroblasts (Supplementary figure 1).

The differentiation potential of patient fibroblasts compared to controls was assessed by quantitative analysis of SMA positive cells after transdifferentiation (Figure 3A). Western blotting showed a SMA expression below the 1xSD range of the controls in FB#1 p.R287W and FB#2 p.R287W. In contrast, the SMA expression in FB#3 p.F248Pfs*62 was significantly increased compared to all other cell lines (p=<0.0001-0.0097) (Figure 3B and 3C). SMA expression in VUS FB#5 p.I396T was slightly above the 1x SD range, but do not significantly differ from controls (p=0.9968) (Figure 3B and 3C). Furthermore, the Western blots indicate that SM22 expression is reduced in FB#1 p.R287W, FB#2 p.R287W and FB#4 p.R268C, although this reduction is not significant (Figure 3B and 3D). In contrast, expression of SM22 in FB#3 p.F248Pfs*62 was significantly increased (Figure 3B and 3D).

**Figure 3.**
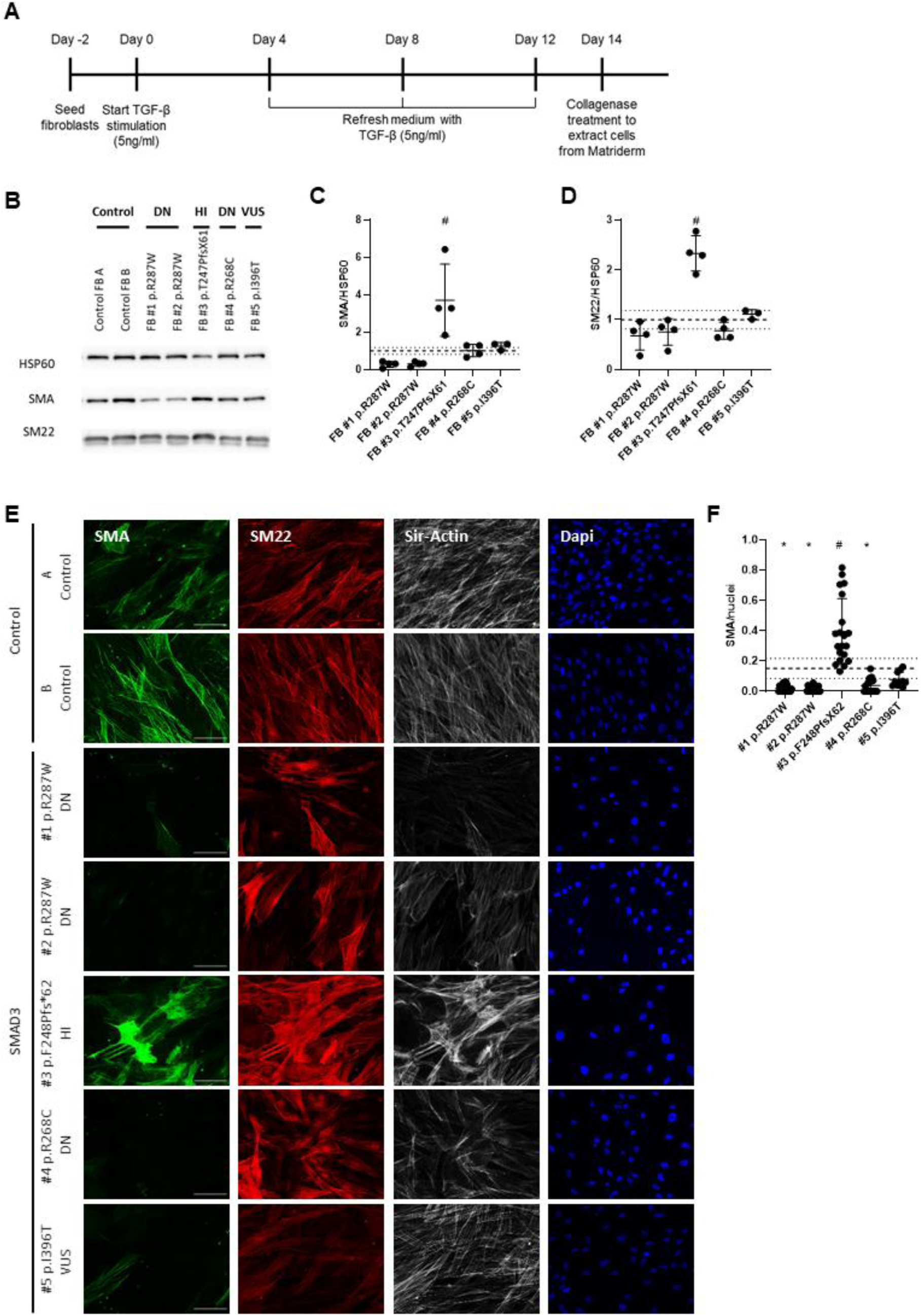
Transdifferentiation potential of SMAD3 fibroblasts is reduced. **A)** Time line of TGF-β induced transdifferentiation of fibroblasts. **B)** Western blots detecting SMA and SM22 in transdifferentiated fibroblasts. HSP60 levels serve as a loading control. **C)** Quantification of SMA levels in B. **D)** Quantification of SM22 levels in D. **E)** Immunofluorescent staining of SMA (green), SM22 (red), F-actin (gray) and DAPI (blue) after 14 days of transdifferentiation of SMAD3 fibroblasts and controls. Scale bar represents 100 µm. **F)** Quantification of SMA positive cells divided by the total number of nuclei. Grey dashed line represents the mean of the controls and black dotted lines represent 1x SD range of the controls. The 1xSD range represents the normal variation of controls. * significantly decreased compared to controls # significantly increased compared to controls and other SMAD3 cell lines

FB#1 p.R287W, FB#2 p.R287W and FB#4 p.R268C myofibroblasts visually showed a reduction of SMA expression in immunofluorescence (Figure 3E). This reduction was confirmed by quantification (Figure 3F), with a mean SMA expression below the 1x SD range (p<0.0001-0.0003). The decrease in SMA expression of VUS FB#5 p.I396T was not significant (p=0.3066). However, since the SMA expression of the VUS is below the 1x SD, the variant is not expected to be HI. On the other hand, FB#3 p.F248Pfs*62 showed a significant increase in SMA expression compared to controls (p<0.0001) as well as to the other *SMAD3* cell lines (p<0.0001). Quantification of SM22 IF staining did not show significant differences (data not shown). Of note, fibroblasts that were not stimulated by TGF-β showed almost no SMA expression and low SM22 expression (data not shown).

In conclusion for this dataset, SMA and SM22 Western blots and immunofluorescence staining indicate a decreased differentiation potential in DN cell lines FB#1 p.R287W, FB#2 p.R287W and FB#4 p.R268C. In contrast, HI cell line FB#3 p.F248Pfs*62 showed an increased differentiation potential. VUS cell line FB#5 p.I396T also showed a reduction of SMA expression assessed by IF staining, although less pronounced compared to the pathogenic DN SMAD3 variants. In addition, the expression of contractile marker MYH11 and synthetic marker vimentin in myofibroblasts was analyzed (Supplemental figure 2), which did not show significant differences.

#### Differentiated DN SMAD3 myofibroblasts show decreased fibrillin-1 ECM deposition

We examined the fibrillin-1 ECM deposition of myofibroblasts from *SMAD3* patients and controls after 7 days of culture (Figure 4).

**Figure 4.**
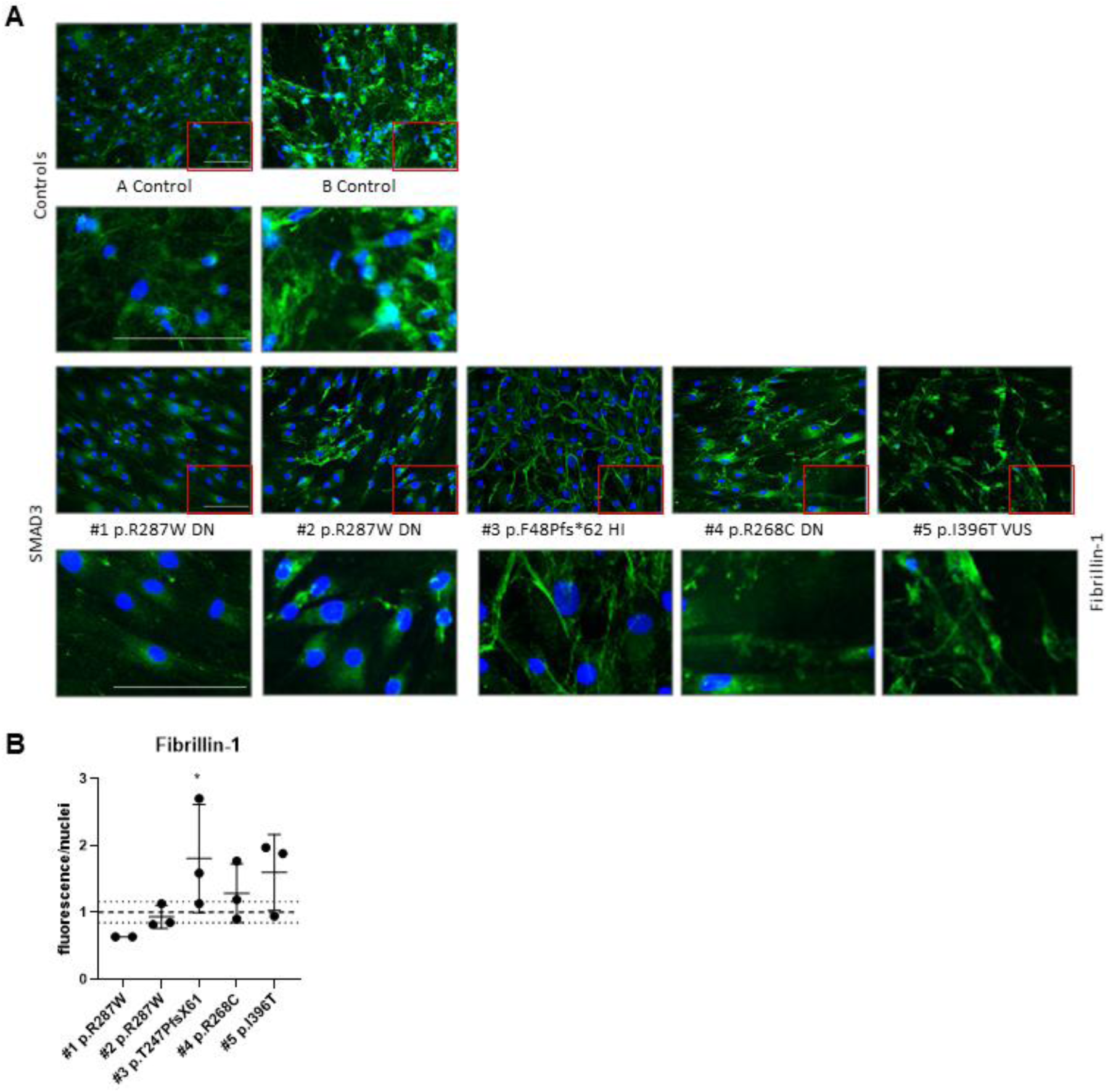
ECM proteins in transdifferentiated fibroblasts. **A)** Fibrillin-1 staining after 7 days of culture. **B)** Quantification of Fibrillin-1 staining. Scale bar represents 100 µm. * significantly increased compared to controls

All cells did show some expression of fibrillin-1, however the DN cell lines show less elongated fibers compared to the controls and HI cell lines.

### Functional assays in vascular smooth muscle cells

#### An HI SMAD3 P/LP variant promotes the expression of contractile marker SMA

To examine whether VSMCs behave similarly as myofibroblasts, VSMCs of *SMAD3* patients and controls were characterized by staining for contractile and synthetic VSMC markers (Figure 5). The expression of contractile marker SMA in HI *SMAD3* VSMC #7 clearly exceeded the 1x SD range of controls on Western blot and was significantly increased compared to controls and DN *SMAD3* VSMCs (Figure 5A-5B). This increased SMA expression was confirmed by IF (Figure 5D and 5E). The SMA expression of the DN *SMAD3* VSMCs was within the 1x SD range and did not significantly differ from controls (Figure 5D and 5E). Although SM22 expression of HI VSMC #7 exceeded the 1x SD range for controls on Western blot, this difference was not significant (Figure 5A and 5C). In addition, quantification of the SM22 staining did not show significant differences (data not shown). Furthermore, Western blot did not show clear differences in vimentin expression, a marker for synthetic VSMCs (Supplemental figure 3).

**Figure 5.**
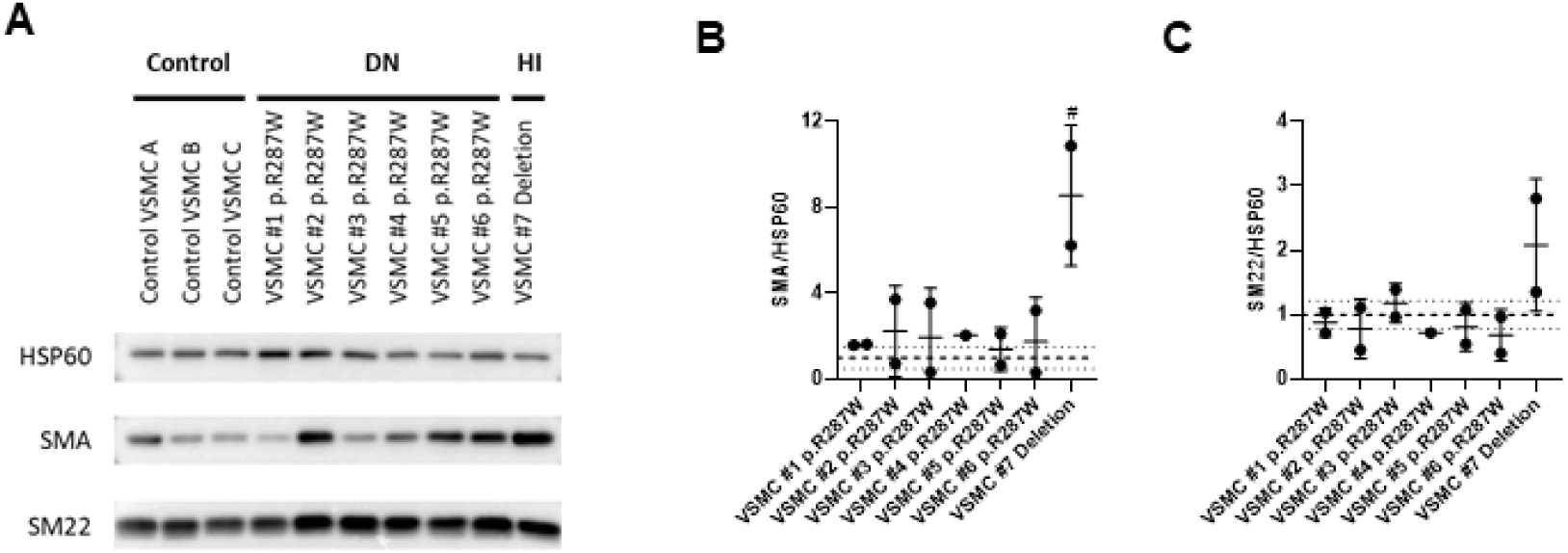
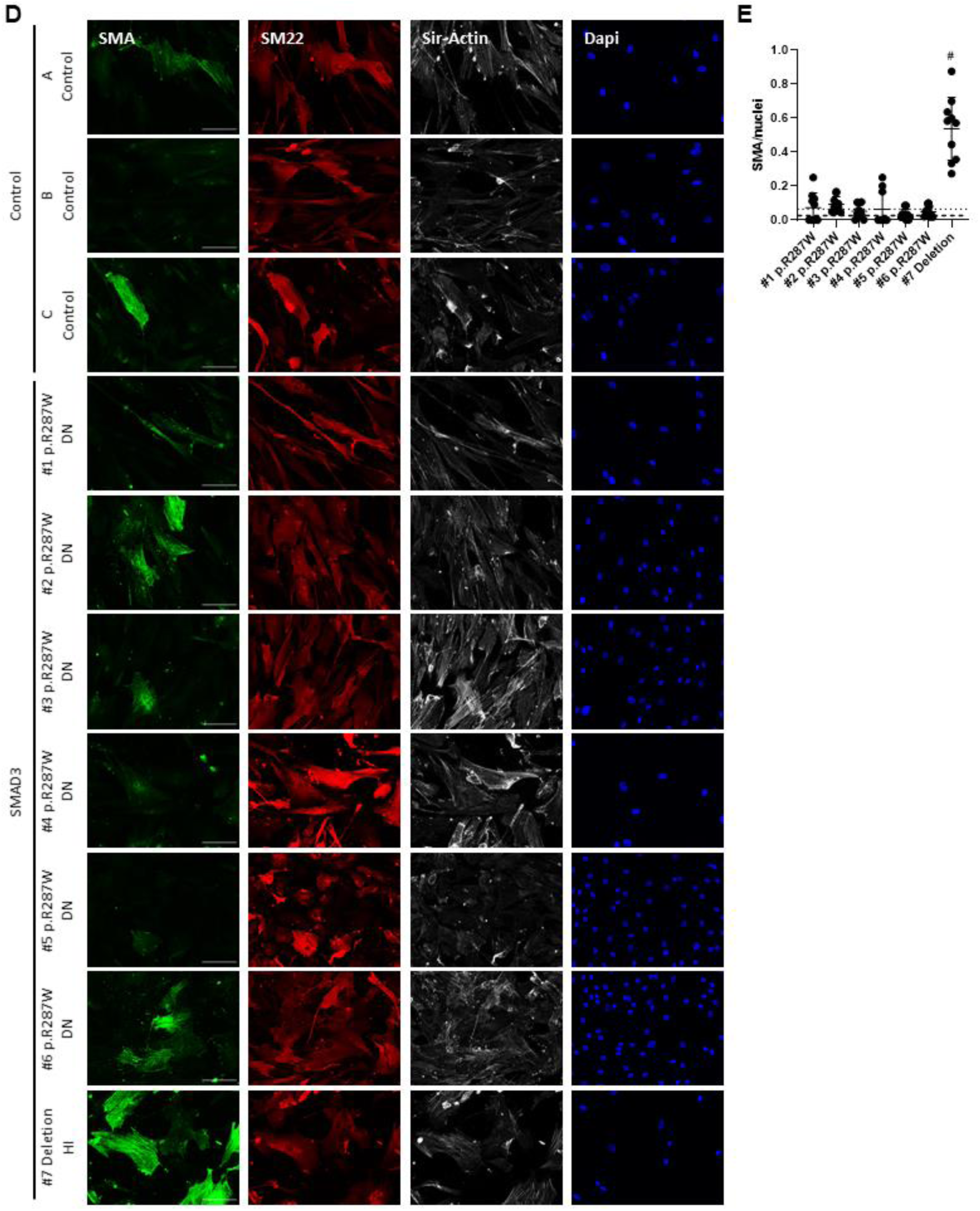
VSMC characterization. **A)** Western blots detecting SMA and SM22 in VSMCs. HSP60 levels serve as a loading control. **B)** Quantification of SMA levels in A. **C)** Quantification of SM22 levels in A. **D)** Immunofluorescent staining of SMA (green), SM22 (red), F-actin (gray) and DAPI (blue) of SMAD3 VSMCs and controls. Scale bar represents 100 µm. **E)** Quantification of SMA positive cells divided by the total number of nuclei. Grey dashed line represents the mean of the controls and black dotted lines represent 1x SD range of the controls. # significantly increased compared to controls and other SMAD3 cell lines.

### A HI *SMAD3* P/LP variant promotes the expression of different *MYH11* isoforms in VSMCs

Another marker for contractile VSMCs is MYH11. Western blot showed increased MYH11 expression for HI *SMAD3* VSMC #7 compared to controls and DN *SMAD3* VSMCs (Figure 6A), which was confirmed with IF (Figure 6B). The expression of *MYH11* in DN *SMAD3* VSMCs (VSMC #1-6) was similar to controls. DNA sequencing of *MYH11* was performed to examine whether the increased MYH11 expression was caused by a *MYH11* P/LP variant. However, no alterations in *MYH11* were observed. Interestingly, the increased MYH11 expression in HI VSMC #7 seems to be caused by expression of a larger MYH11 protein isoform. Different isoforms of *MYH11*, caused by alternative splicing, are known to be expressed during embryonal development (39) and are indicated in Figure 6C. A PCR analysis of cDNA prepared from control VSMC A and VSMC #7 indicated different expression levels of the *MYH11* isoforms in VSMC #7, with highest expression of isoform SM1A (Figure 6D, Table 2). This was confirmed by splice analysis on RNA sequencing data in VSMC #7 and VSMC A, revealing alternative splicing of exon 5b (5’ end) and the exon generating the short carboxyl terminus (3’ end) in VSMC #7 (Figure 6E). Based on the number of alternative splicing events causing exclusion of both exons, isoform SM1A is expected to be most expressed (Figure 6F). Expression of isoform SM1A results in a protein with a length of 1972 amino acids, which is the second largest isoform of MYH11. This can explain the additional band observed on Western blot indicating the expression of a larger MYH11 isoform compared to control VSMCs.

**Figure 6.**
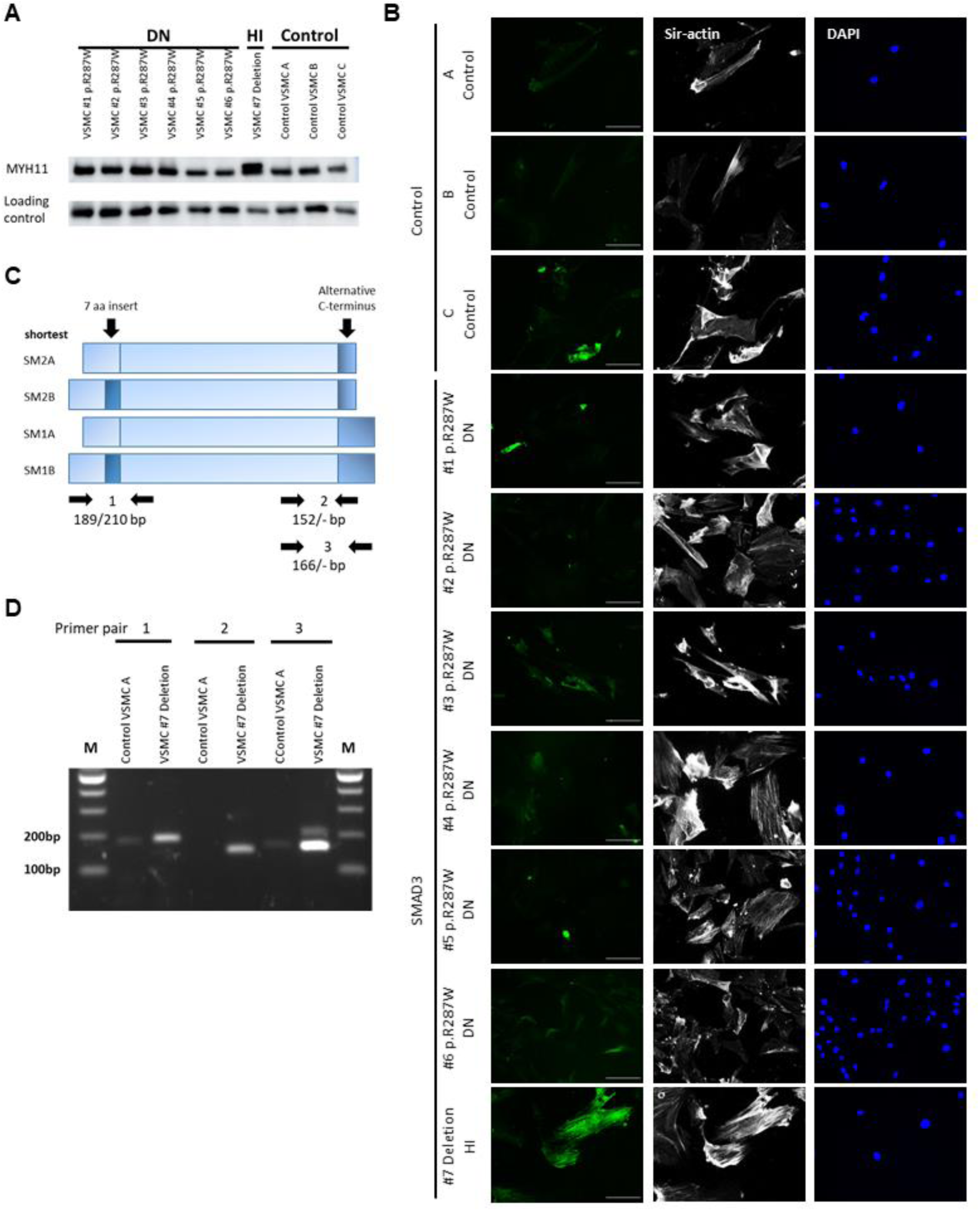
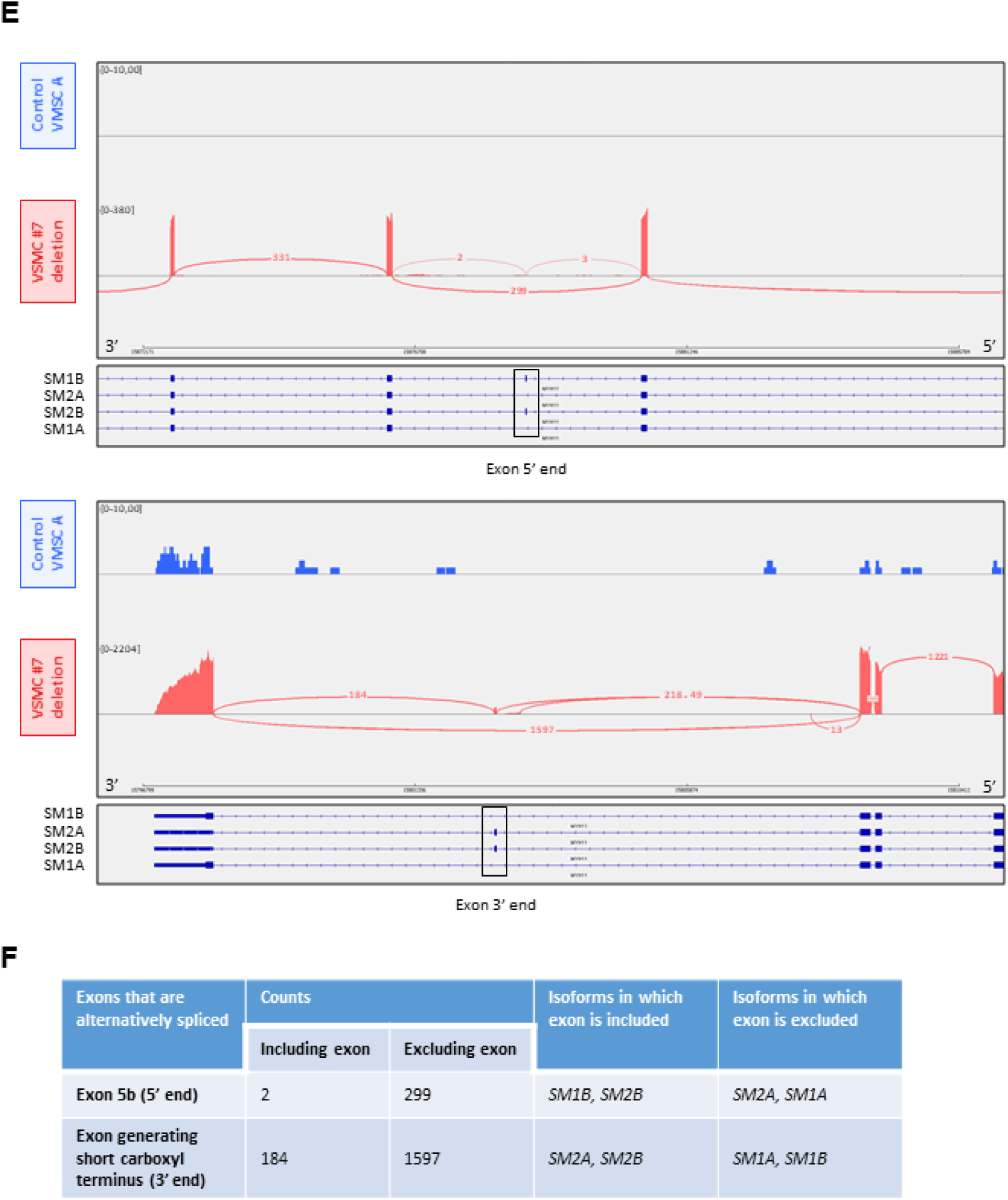
MYH11 expression in VSMCs. **A)** Western blot detecting MYH11 in VSMCs. Β-catenin levels serve as a loading control. **B)** Immunofluorescent staining of MYH11 (green), F-actin (gray) and DAPI (blue) of SMAD3 VSMCs and controls. Scale bar represents 100 µm. **C)** Schematic representation of different MYH11 isoforms caused by alternative splicing. Indicated the primer pairs to perform PCRs on cDNA to detect different isoforms **D)** PCR on cDNA to detect different isoforms of MYH11 in Control VSMC A and SMAD3 VSMC #7 Deletion. Table 4 gives an overview of expected sizes of PCR products for the different isoforms. **E)** Sashimi plot for the alternatively spliced exons on the 5’ end and 3’ end of the MYH11 gene in patient VSMCs (#7 deletion) and control VSMCs. Per-base expression is plotted on the y-axis of the Sashimi plot and genomic coordinates on the x-axis. The four different isoforms of MYH11 (SM1B, SM2A, SM2B and SM1A) are indicated below the Sashimi plot. Alternative splicing events of both exons are detected in the patient VSMCs, whereas none are detected in the control VSMCs. **F)** Table showing the number of splice events resulting in either inclusion or exclusion of the exon on the 5’ end and the exon on the 3’ end of MYH11. Since the largest number of splice events result in exclusion of both the exon on the 5’ end and the exon on the 3’ end, isoform SM1A is expected to be most expressed.

**Table 4.** Overview characteristics transdifferentiated fibroblasts and VSMCs compared to controls.

|  | Expression VSMC markers |  |  |  |  |  |  |  | ECM formation |  |  |  |
| --- | --- | --- | --- | --- | --- | --- | --- | --- | --- | --- | --- | --- |
|  | Western blot |  |  |  | Immunofluorescence |  |  |  | ECM staining |  |  |  |
|  | SMA | SM22 | MYH11 | Vimentin | SMA | SM22 | MYH11 | Vimentin | Fibronectin | Fibrillin-1 | Fibulin-4 | Fibulin-5 |
| **TD fibroblasts** |  |  |  |  |  |  |  |  |  |  |  |  |
| DN | ↓* | ↓ | ~ | ~ | ↓* | ↓ | ~ | ~ | ↓ | ↓ | ↓ | ↓ |
| HI | ↑* | ↑* | ~ | ~ | ↑* | ↑ | ~ | ~ | ↑ | ~ | ~↑ | ~ |
| **VSMCs** |  |  |  |  |  |  |  |  |  |  |  |  |
| DN | ~ | ~ | ~ | ~ | ~ | ~ | ~ | ~ | ~ | ~ | ~ | ~ |
| HI | ↑* | ↑ | ↑* | ~ | ↑ | ↑ | ↑ | ~ | ~ | ~ | ~ | ~ |

Summary of functional assays in transdifferentiated fibroblasts and VSMCs of individuals with a P/LP SMAD3 variant. TD; transdifferentiated, VSMCs; vascular smooth muscle cells, SMA; smooth muscle actin, SM22; smooth muscle 22, ECM; extra cellular matrix. Significant differences are visualized with \*. No differences compared to controls is visualized with ~.

#### SMAD3 mutated VSMCs show delayed fibrillin-1 deposition

We examined the deposition of ECM component fibrillin-1 of VSMCs from *SMAD3* patients and controls after 7 days (Figure 7) and after 14 days (Supplemental figure 4).

**Figure 7.**
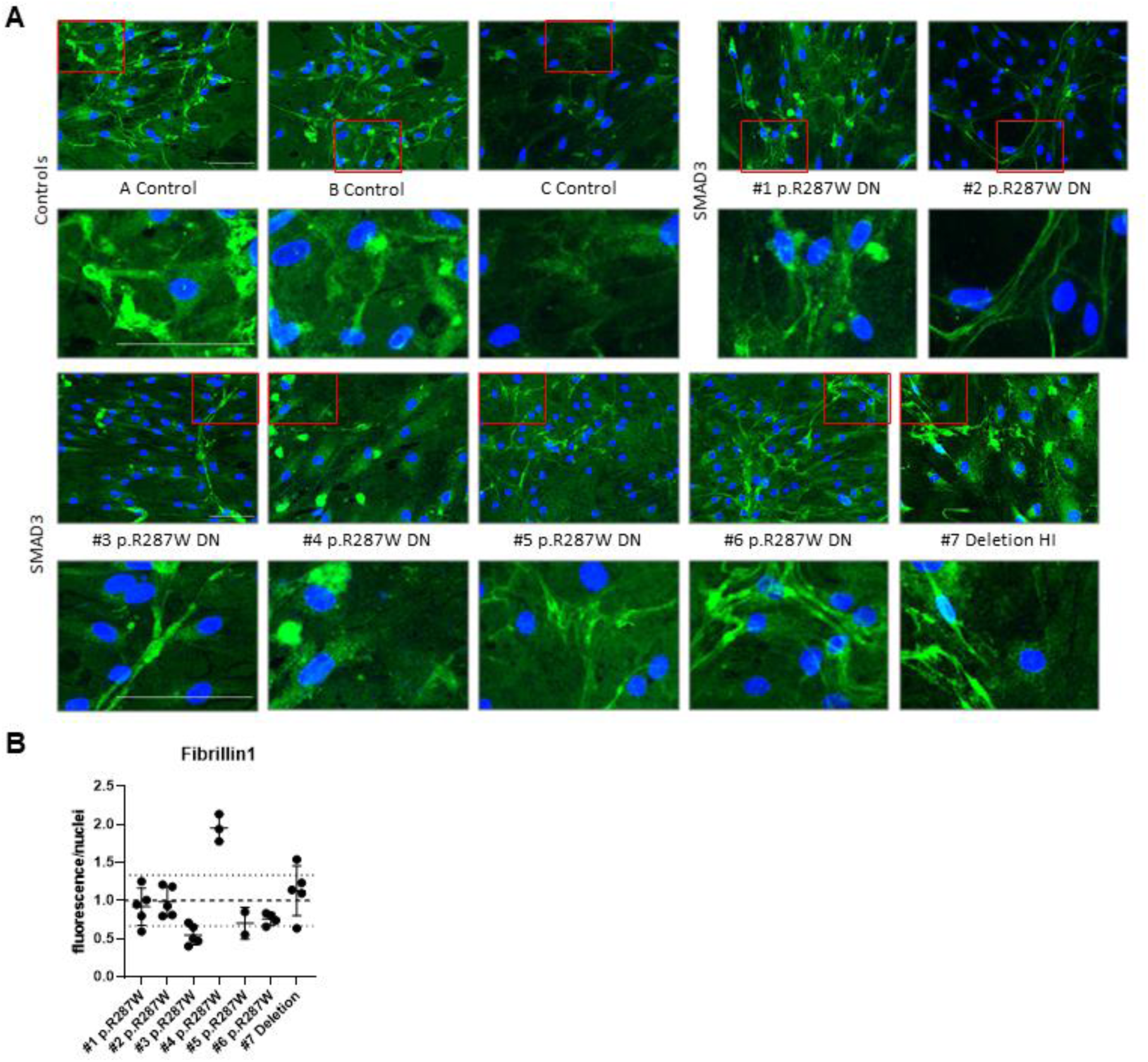
ECM proteins in VMSC. Immunofluorescent images show deposition of ECM components after 7 days of culture. Scale bar represents 100 µm. **A)** Fibrillin-1 staining. **B)** Quantification of Fibrillin-1 staining.

Fibrillin-1 staining was mainly reduced in *SMAD3* VSMC #3, #5, #6 p.R287W after 7 days (Figure 7) and VSMC #4 seems to form more fibrillin-1. After 14 days, all cell lines show comparable ECM deposition (Supplemental figure 4B).

## DISCUSSION

Functional assays on patient-derived differentiated fibroblasts and VSMCs with a P/LP variant in *SMAD3* were performed in order to elucidate the molecular and cellular effects of these *SMAD3* variants, to improve interpretation of VUS. There is an unmet need for a fast and reliable readout in the diagnostic setting, since the number of identified VUS is growing and adequate interpretation of these VUS is essential for patient management. Besides, the comparison of the assays of these two cell types will provide further insight into the reliability of using differentiation of fibroblasts instead of VSMCs, as patient fibroblasts are more easily accessible. A summary of the molecular and cellular characteristics established in this study is provided in Table 4.

1. The differentiation potential of myofibroblasts differed between different types of *SMAD3* variants. A strong reduction is seen in SMA expression after differentiation into myofibroblasts in the DN *SMAD3* myofibroblasts, which is in line with previous results obtained for FB#1 p.R287W (40). In contrast to the DN *SMAD3* myofibroblasts, HI *SMAD3* myofibroblasts show an increased differentiation potential. Hence, not all *SMAD3* variants affect differentiation into myofibroblasts after TGF-β stimulation to the same extent, which might point to a different underlying molecular disease mechanism for DN and HI *SMAD3* variants.
The expression of contractile markers SMA and SM22 was increased in HI *SMAD3* VSMCs, while there is no significant difference in DN *SMAD3* VSMCs compared to controls. The synthetic marker vimentin is not reduced in those cells, indicating that despite an increase in contractile markers there is no clear shift towards the contractile phenotype.
2. The ECM formation is less structured in DN *SMAD3* myofibroblasts for fibrillin-1 compared to HI *SMAD3* myofibroblasts. This finding is interestingly, since P/LP variants in genes involved in formation and integrity of the ECM, like Fibrillin-1, generally lead to ECM accumulation. Furthermore, ECM components are still formed in VSMCs, but the formation seems to be delayed in some DN *SMAD3* VSMCs. This is in line with previous findings for Smad3 mouse models (41–43). The reduced ECM formation is likely caused by dysregulation of the downstream TGF-β pathway. This dysregulation reduces downstream transcription of genes, including ECM components and matrix metalloproteases (MMPs), which are important for matrix remodeling. The reduced transcription of ECM components and MMPs disrupts sufficient ECM formation and weakens the aortic structure.
3. SMAD2 and SMAD3 phosphorylation was not altered in LDS3 patient-derived fibroblasts after TGF-β stimulation. This indicates that initial activation of the TGF-β pathway can still occur in cell lines with P/LP *SMAD3* variants with either DN or HI effect. The antibodies for SMAD3 and pSMAD3 recognize the part of SMAD3 where the P/LP variants are not located and the P/LP variants are not expected to alter the phosphorylation sites. Therefore, the absence in different phosphorylation between healthy and mutated SMAD3 will not be caused by the action of the antibodies.

### Comparison of functional assays between myofibroblasts and VSMCs

We compared the differentiation potential and ECM formation between fibroblasts differentiated into myofibroblasts and VSMCs, to examine whether myofibroblasts are an alternative model to study the molecular mechanism of *SMAD3* variants.

The SMA and SM22 expression patterns were equal for HI *SMAD3* myofibroblasts and VSMCs. In contrast, SMA and SM22 expression was reduced in DN *SMAD3* myofibroblasts compared to controls and not in VSMCs. Besides, myofibroblasts showed a reduced elongated fibrillin-1 fiber formation in DN *SMAD3* cell lines compared to controls, whereas no differences in ECM formation were observed in either DN *SMAD3* VSMCs or HI *SMAD3* VSMCs compared to controls.

The different origin of fibroblasts and VSMCs might explain the different outcomes in SMA and SM22 expression and ECM formation between these cell types. VSMCs are isolated from different parts of the aorta, while fibroblasts are all obtained from the inner side of the upper arm. Additionally, it is important to realize that the embryonic origin can vary between VSMCs, since the aorta is derived from distinct embryonic sources. The ascending aorta is derived from neural crest (NC), whereas the descending aorta originates from somatic mesoderm (44). Several studies suggest that individual VSMC characteristics, including gene expression, are determined by the embryonic origin (45–47). Besides the embryonic origin, the region of the aorta can contribute to the VSMC characteristics, since previous studies have shown that the VSMC characteristics differ based on the location in the vascular tree (47). Furthermore, VSMCs play an important role in arterial remodelling to maintain arterial structure and function (48). Due to biomechanical and biochemical stressors, VSMCs will lose contractile markers and differentiate to a synthetic VSMC phenotype to induce proliferation and migration (49). The different embryonic origins of VSMCs and phenotypic switch under biological stress signals might influence the results not only between fibroblasts and VSMCs but also between different VSMCs lines. Therefore, large numbers of VSMCs are needed to examine whether specific results are based on phenotypic switches or due to a genetic variant. In conclusion, functional assays for differentiation potential and ECM formation can distinguish between DN and HI *SMAD3* variants. Differentiation of fibroblasts seems to be a more suitable method for these functional assays as an alternative for VSMCs.

### Differences between HI and DN *SMAD3* variants

The expression of VSMC markers showed opposing results for DN and HI *SMAD3* mutant cells; SMA and SM22 expression is reduced in DN *SMAD3* variants and increased in HI *SMAD3* variants. This suggests that there might be a different effect on the TGF-β and bone morphogenetic protein (BMP) signalling within the cells. BMP signaling plays an important role in vascular remodeling, maintenance of joint integrity, the initiation of fracture repair, among others (50). The TGF-β pathway signaling can occur via a canonical or non-canonical route. The canonical route includes two pathways, namely via TGF-β ligands and via BMP ligands. If the TGF-β canonical pathway is activated by TGF-β ligands, SMAD4 forms complexes with SMAD2/3. If activated by BMP ligands, SMAD4 forms complexes with SMAD1/5/8 (R-SMADs). We speculate about the role of SMAD4 in the different effects of DN and HI variants, since SMAD4 is present in the canonical TGF-β signaling and the BMP signaling pathway.

The phosphorylation of SMAD1/5/8 is indirectly induced by BMPs. Phosphorylated SMAD1/5/8 associates with SMAD4 and regulates gene expression in the cell nucleus. If more SMAD4 is available, more complexes with SMAD1/5/8 will be formed resulting in more gene regulation via the BMP signaling. Since only half of the SMAD3 protein is available in cells with HI SMAD3 variants, a reduced amount of complexes with SMAD4 will be formed. This will result in less ‘trapped’ SMAD4 and thus a larger amount of freely available SMAD4 to form complexes with SMAD1/5/8 and regulate downstream genes, including *SMA*. The increased regulation of downstream genes might explain the increased differentiation potential in HI *SMAD3* mutated cells compared to controls and DN *SMAD3* mutated cells.

### *MYH11* isoforms

A strongly increased *MYH11* expression was only noticed in the HI *SMAD3* VSMC and seems to be caused by the expression of an additional protein isoform. Four different isoforms are known for MYH11; the expression of which differs during embryonic development and varies between tissue types (39). The SM1 isoforms are mostly present in the embryonic stage, whereas the SM2/SM1 ratio increases during cell maturation. PCRs and RNA sequencing confirmed the altered expression of *MYH11* isoforms in the HI *SMAD3* VSMC. The SM1A isoform that is normally present during embryonic development is probably most abundant. We hypothesize that the altered expression of MYH11 isoforms might be a result of altered *SMAD3* expression, since these HI VMSCs did potentially undergo a different maturation/differentiation process compared to controls and to *SMAD3* cell lines with a DN P/LP variant. Another hypothesis is based on the knowledge that tissues can revert to embryonic characteristics during stress. Reverting to the embryonic developmental program gives cells of a specific organ the ability to repair and maintain function, which is previously shown in for example heart tissue (51, 52). Due to the presence of a HI *SMAD3* P/LP variant, these VSMCs have endured stress and might be reverted to their embryonic developmental program. The HI myofibroblast cell line did not show altered expression of *MYH11*, which is possibly caused by the very low *MYH11* expression in fibroblasts (53). As we only had VMSCs of one HI *SMAD3* patient, we were not able to reproduce these data in independent HI *SMAD3* VSMC lines.

### Correlation between phenotypic variability and VSMC markers

Phenotype-genotype correlations between HI and DN P/LP *SMAD3* variants are previously studied by Hostetler et al. in 212 individuals with a P/LP *SMAD3* variant (31). Individuals with a DN *SMAD3* variant in the MH2 domain were significantly younger at first aortic event compared to those with a DN variant in the MH1 domain or a HI variant. Our retrospective analysis in *SMAD3* individuals seems to confirm the observations of Hostetler et al. (31). The phenotypic variability between DN and HI variants might in part be explained by the different underlying molecular mechanisms like the observed opposite expression of VSMC markers between DN and HI myofibroblasts.

Unfortunately, we were not able to determine the pathogenicity of the VUS based on the differentiation potential and ECM formation. The characteristics of the VUS fibroblasts were intermediate between those of the controls and P/LP DN *SMAD3* variants and it is not possible to draw conclusions based on these small differences with the limited amount of available cell lines in this study. Therefore, an extension of this research including a larger number of cell lines is necessary to further investigate the SMA and SM22 expression and ECM formation in P/LP SMAD3 mutant cell lines. Moreover, it would be interesting to investigate whether the observed differences in SMA and SM22 expression and ECM formation is specific for P/LP *SMAD3* variants or whether these features are observed in other aneurysm genes as well.

## ACKNOWLEDGEMENTS

We would like to thank Bianca de Graaf and Joyce Burger for isolating VSMC cell lines and Joyce Lebbink for analyzing the effects of variants on protein structure and the department of Clinical Genetics, laboratory diagnostics, for the preparation and storage of patient fibroblasts.

